# Disclosing the actual efficiency of G-quadruplex-DNA–disrupting small molecules

**DOI:** 10.1101/2020.11.16.382176

**Authors:** Jérémie Mitteaux, Pauline Lejault, Marc Pirrotta, Filip Wojciechowski, Alexandra Joubert, Nicolas Desbois, Claude P. Gros, Robert H. E. Hudson, Jean-Baptiste Boulé, Anton Granzhan, David Monchaud

## Abstract

The quest for small molecules that avidly bind to G-quadruplex-DNA (G4-DNA, or G4), so called G4-ligands, has invigorated the G4 research field from its very inception. Massive efforts have been invested to *i*- screen or design G4-ligands, *ii*- evaluate their G4-interacting properties *in vitro* through a series of now widely accepted and routinely implemented assays, and *iii*- use them as unique chemical biology tools to interrogate cellular networks that might involve G4s. In sharp contrast, only uncoordinated efforts at developing small molecules aimed at destabilizing G4s have been invested to date, even though it is now recognized that such molecular tools would have tremendous application to neurobiology as many genetic and age-related diseases are caused by an over-representation of G4s, itself caused by a deficiency of G4-resolving enzymes, the G4-helicases. Herein, we report on our double effort to *i*- develop a reliable *in vitro* assay to identify molecules able to destabilize G4s, the G4-unfold assay, and *ii*- fully characterize the first prototype of G4-disrupting small molecule, a phenylpyrrolcytosine (PhpC)-based G-clamp analog.

## Introduction

The existence of a higher-order, four-stranded DNA structure known as G-quadruplex-DNA (G4-DNA, or G4)^[1]^ within functional human cells is now established through different technologies, including structure-specific isolation and identification techniques (either low-^[2]^ or high-throughput techniques)^[3]^ and optical imaging.^[1c, 4]^ The cellular role of G4s defies easy understanding and facile explanations as G4s fold from numerous regions of the human genome (>500,000),^[1b, 5]^ not systematically committed to key regulatory functions. However, their formation is unquestionably coupled with DNA transactions (transcription and replication)^[6]^ as a result of both duplex melting and supercoiling originating in the motion of DNA/RNA polymerases along the duplex stem. This makes G4 formation a possible impediment to DNA transactions as G4s possess high thermodynamic stablity and may be persistent in genomic DNA, therefore representing solid physical obstacles to helicase and polymerase processivity.

To prevent such a situation, DNA/RNA polymerases coordinate their action with enzymes that unwind G4s, known as G4-helicases.^[7]^ The uncoupling of the polymerase and helicase activity creates a situation of crisis that ultimately leads to DNA damage and genome instability. This might arise as a result of either an abnormal G4 stabilization (for instance, *via* externally added G4-stabilizing compounds, or G4-ligands) or a G4-helicase impairment. This later hypothesis is now well documented as the loss-of-function mutation of helicases is linked to severe genetic conditions. Numerous human G4-helicases have been characterized, belonging to two different helicase super-families (SF1 and SF2),^[8]^ which include the SF1 Pif1^[9]^ and the SF2 BLM and^[10]^ WRN^[10b, 11]^ (RecQ-like helicase subfamily) as well as FANCJ^[12]^ and DDX1^[13]^ (Fe-S helicase subfamily). Each of these enzymes is associated to a human genetic disease: the Bloom syndrome (growth retardation, immunodeficiency) is caused by a mutation of BLM,^[14]^ the Werner syndrome (adult progeria) by that of WRN,^[15]^ the Fanconi anemia (developmental abnormalities, bone marrow failure) by FANCJ,^[16]^ and the Warsaw Breakage syndrome (impaired growth, intellectual disability) by DDX1.^[13a, 17]^ Pif1 deficiency is more generally associated with cancer predisposition.^[18]^ Also, we recently showed that an over-representation of G4s originating in either age-associated changes in the activity of G4-modulating proteins and in G4 stabilization by G4-ligands may accelerate brain aging and foster neurological disorders.^[19]^

In light of this, it is surprising that most, if not all, chemical biology efforts have been invested in the quest of chemicals that stabilize G4s (G4-ligands, with >1,500 PubMed entries to date)^[20]^ rather than compounds capable of unfolding G4s to rescue helicase impairment. Beyond historical reasons (the first therapy-oriented G4-ligand was reported in 1997 to stabilize telomeric G4 in order to impair the cancer-relevant telomerase enzymatic complex),^[21]^ G4 stabilization can be a strategically wise way to inflict severe damage to the genome of cancer cells, the selectivity of the treatment relying on their flawed repertoire of DNA damage signaling and repair capabilities (collectively known as DNA damage response, or DDR) as compared to healthy cells.^[22]^ However, this does not explain the paucity of validated prototypes of G4 unwinders.

Over the past years, some examples of G4-unwinders have been reported,^[23]^ such as: *i*- the porphyrin TMPyP4,^§^ shown to unfold G4-DNA that folds from d[(CGG)_n_] trinucleotide repeats, whose expansion is involved in the Fragile X syndrome,^[24]^ and from the d[(GGGGCC)_n_] hexanucleotide repeats, whose expansion is linked to ALS/FTD;^[25]^ *ii*- an anthrathiophedione derivative, shown to unfold the human telomeric G4-forming sequence d[(TTAGGG)_n_];^[26]^ *iii*- the triarylpyridine TAP1, reported to disrupt the G4 that folds from a sequence of the c-kit promoter;^[27]^ *iv*- a series of stiff-stilbenes found to regulate the folding/unfolding of the telomeric G4 in a photoresponsive manner;^[28]^ along with copper ion and complexes,^[29]^ urea^[30]^ and natural polyamines (*e.g*., spermine).^[31]^ There is, however, not broad consensus on the use of these chemicals as surrogates for helicases, given that no in-depth cellular investigations have been yet performed. This might be due to the doubts about their actual unfolding activity: TMPyP4 is an illustrative example of this conundrum, as it was studied for more than two decades as a G4-stabilizer^[20, 32]^ and is now reported as a G4-unfolder.^[24–25, 33]^ These surprising results are however in line with previous reports in which the paradoxical behavior of porphyrin derivatives is described, either as a function of the nature of their metallic complexes (the platinum complex of TMPyP4 destabilizes G4s while the free base TMPyP4 and both the zinc and copper complexes stabilize them)^[34]^ or as a function of the techniques implemented (the spermine-decorated porphyrin TCPPSpm4^§^ can destabilize or stabilize G4s if gradual or blunt additions are performed, respectively).^[35]^

We reasoned that a possible problem is the lack of reliable *in vitro* assay to assess the G4 unfolding properties of chemicals. Dozens of assays have been developed to quantify the G4-intercating/stabilizing properties of ligands;^[36]^ in sharp contrast, no reliable, systematic and high-throughput screening (HTS) assay is available for studying G4 disruption: the examples described above have been documented *via* different assays (chiefly polyacrylamide gel electrophoresis (PAGE) and circular dichroism (CD), *vide infra*) without systematic comparison and analysis of *ad hoc* controls.

### G4-unfold design

To tackle this issue, we decided to revisit the fluorescence-based helicase assay developed by Mendoza, Bourdoncle and coworkers in which the G4 opening ability of the helicase Pif1 was quantified *via* a HTS-compatible fluorescence analysis (Figure 1A).^[37]^ To this end, the authors designed a bimolecular DNA system named S-htelo comprising a 49-nt long oligodeoxynucleotide (ODN), which includes a 5’ d[5’(A)_11_3’] tail for Pif1 loading, the human telomeric sequence d[5’(G_3_T_2_A)_3_G_3_3’] and a 17-nt 3’ tail labeled with dabcyl (consequently named dabcyl-labelled 49-nt ODN), and a 15-nt long ODN that is complementary to the 3’ tail of S-htelo and labeled with FAM on its 5’-end (consequently named FAM-labelled 15-nt ODN). When hybridized, this system possesses a single-stranded region (for helicase loading), a folded G4 and a duplex region that ends with a FRET pair in which the FAM fluorescence is quenched by the proximal dabcyl. The helicase assay *per se* is triggered by the addition of both Pif1 (0.5 mol. equiv.), which, in presence of an excess of ATP (4.5 mM), unfolds the system in a 5’-to-3’ manner. The strand separation is then monitored through the enhancement of the FAM fluorescence. The reverse reaction is suppressed by the addition of a 15-nt ODN named Trap (5 mol. equiv.), complementary to the FAM-labelled 15-nt ODN, and the process is driven to completion by the addition of a 49-nt ODN named C-htelo (5 mol. equiv.), fully complementary to dabcyl-labelled 49-nt ODN. This assay was originally developed to quantify Pif1 activity and its inhibition by G4-stabilizing agents (BRACO-19,^[38]^ pyridostatin (PDS),^[39]^ PhenDC3^[40]^ and TrisQ,^[41]^ 25 mol. equiv.).

**Figure 1.**
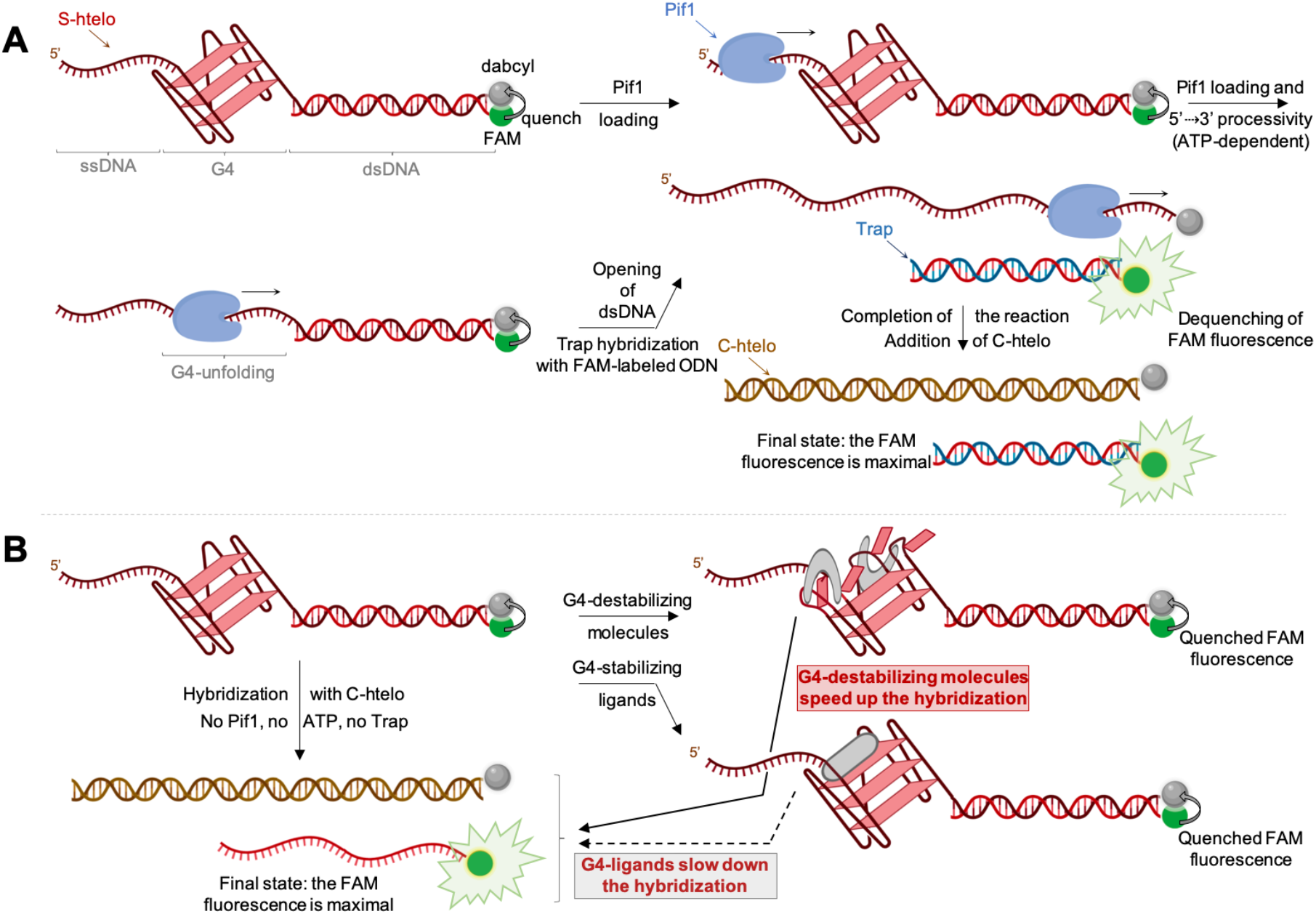
Schematic representation of the helicase assay developed by Mendoza, Bourdoncle *et al*. (**A**) and of the related G4-unfold assay suited to evaluate the G4-destabilizing properties of small molecules (**B**).

This assay is efficient but cannot be conveniently used as a HTS test for screening G4-disrupting molecules because of the limited access to Pif1 helicase that is not commercially available and must be expressed and purified. We reasoned that a modified, simplified version of this assay might be suited to assess the G4-disrupting activity of small molecules: indeed, the kinetics of the final DNA system opening upon addition of C-htelo could be affected by the presence of chemicals, being either slowed down by G4-stabilizing compounds or accelerated by G4-destabilizing compounds. This approach would greatly simplify the protocol, making it a one-step assay in which the initial FAM/dabcyl duplex is incubated with putative candidates whose effect on G4 stability is directly monitored upon addition of C-htelo (no Pif1, no ATP, no Trap). This assay named G4-unfold (Figure 1B) is thus practically convenient, being performed at room temperature for 1h in a 96-well plate format.

### The selection of a representative panel of candidates

To investigate the validity of the G4-unfold assay, we selected a panel of representative compounds (Figure 2) comprising *i*- five porphyrins including the tetracationic TMPyP4^[24–25, 32–33]^ and its PEGylated analogue TEGPy,^§, [42]^ the tetraanionic TPPS^§^ and its PEGylated analogue TArPS,^§,[43]^ and the neutral, water-soluble (PEGylated) TEGP,^§^ to assess the actual efficiency of TMPyP4 and the influence of both the charges (cationic, neutral, anionic) and the PEG arms on the G4-disruption/stabilization ability of porphyrin cores; *ii*- three G4-ligands, PhenDC3,^[40, 44]^ PDS^[39]^ and BRACO19,^[38, 45]^ to calibrate the assay with firmly established G4-stabilizers; *iii*- a TAP1^[27, 46]^ analogue referred to as Terpy; and *iv*- a series of compounds that might be suited to G4-disruption, *i.e*., three macrocyclic bis-naphthalene compounds (or azacyclophanes), 1,5-BisNPO, 2,6-BisNPO and 2,7-BisNPN,^[47]^ whose ability to sandwich and stabilize isolated aromatic compounds (*e.g.*, a wobbling guanine escaping from the external G-quartet upon G4 destabilization) has been demonstrated by both NMR^[48]^ and X-ray crystal structure analysis,^[49]^ and two G-clamp^[50]^ analogues, PhpC and guaPhpC,^§,[51]^ known to strongly interact with Gs thanks to the formation of 4 H-bonds (*versus* 3 for the canonical GC base pair), which could similarly trap and stabilize a flipping guanine.

**Figure 2.**
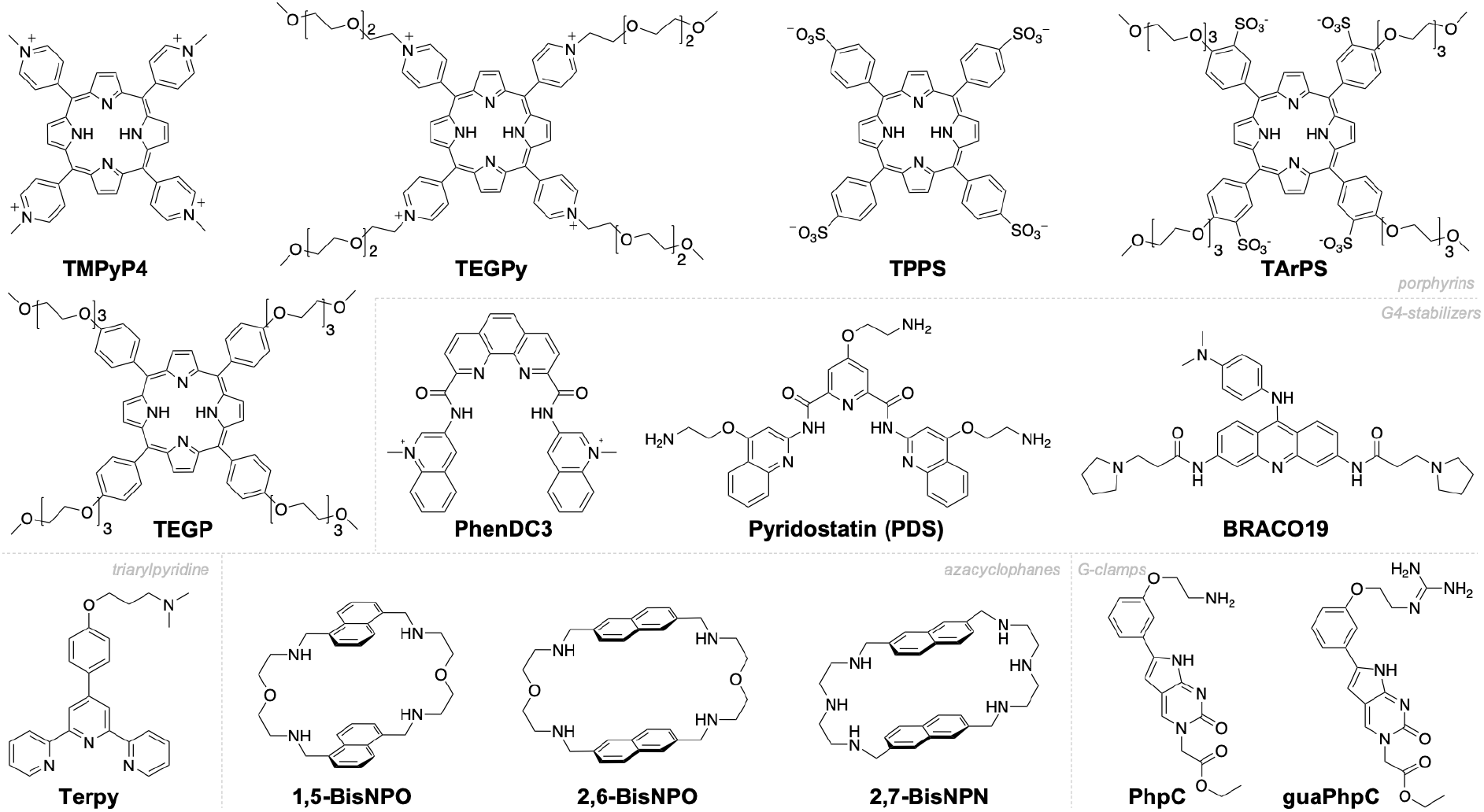
Molecules evaluated in the G4-unfold assay, belonging to the series of porphyrins (TMPyP4, TEGPy, TPPS, TArPS and TEGP), G4-stabilizers (PhenDC3, PDS and BRACO19), triarylpyridines (terpy), azacyclophanes (1,5-BisNPO, 2,6-BisNPO and 2,7-BisNPN) and G-clamp analogues (PhpC and guaPhpC).

### G4-unfold evaluations

First, the kinetics of the FAM/dabcyl duplex opening was decreased by using a lower concentration of C-htelo (2 mol. equiv. *versus* 5 mol. equiv. in the initial setup, with V_0_ = 70.5 *versus* 51.5 s^−1^, respectively) and the effect of chemicals assessed throughout a wider range of concentrations (1, 5, 10 and 20 mol. equiv.). As seen in Figure 3 and in the Supporting Information (Figures S1-6), the presence of the small molecules affects both the kinetics (represented by the slope of the curve after C-hTelo addition) and thermodynamics of the hybridization (represented by the final fluorescence level). This is particularly obvious for the experiments performed with TMPyP4 (Figure 3E), which might originate in several factors (*e.g*., screen effect) that cannot be easily disentangled. A way to circumvent this issue would be to normalize the curves obtained: as seen in Figures S7,8, trends described hereafter are respected when curves are normalized (which removes the thermodynamics contribution), reinforcing our commitment to exploit raw data only, in order to avoid unwarranted data manipulation. The variation of the kinetics of the reaction (n > 4) was comprised between V_0_ = 1.7 and 104.0 s^−1^ and was found strongly dependent on both the nature and the stoichiometry of the chemicals. We selected a 2xS.D. exclusion zone (twice the standard deviation, diagonally hashed zone) around the V_0_ calculated for the reaction without chemical (control (Ctrl), white bar) and ranked the performances of the candidates on the basis of their own 2xS.D. values: compounds that overlap with the exclusion zone of the control appear in grey (affecting the kinetics in a non-significant manner), while those which display higher V_0_ values appear in dark red (accelerating the hybridization) and those with lower V_0_ values in dark cyan (slowing the hybridization).

**Figure 3.**
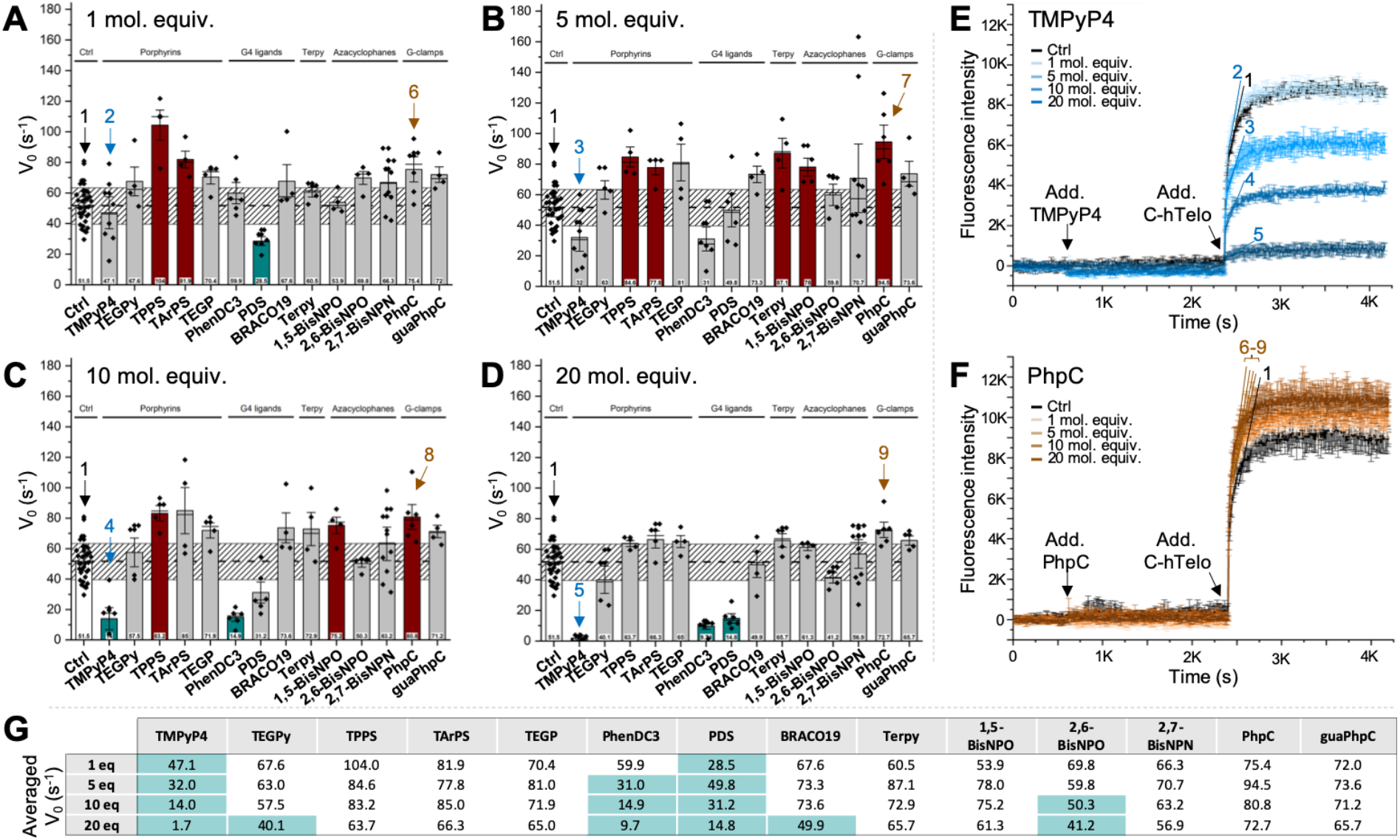
**A-D**. Results (initial velocity V_0_, expressed in s^−1^) of the G4-unfold assay performed with the FAM/dabcyl duplex construct (40 nM) and increasing amounts of 14 compounds (from 1 to 20 mol. equiv.); diamonds are the experimental data (n > 4); bars represent the averaged V_0_; error bars represent standard deviation (S.D.); the diagonally hashed grey zone represent the exclusion zone (calculated as 2xS.D. of the control, Ctrl). **E-F**. Examples of experimental curves (n = 2) obtained with TMPyP4 and PhpC. **G**. Averaged V_0_ values obtained with the G4-unfold assay; values in dark cyan are those below that of the control (V_0_ = 51.5 s^−1^).

Both PDS and PhenDC3 markedly slow down the hybridization (with V_0_ down to 14.8 and 9.7 s^−1^, respectively), thus lending credence to the hypothesis according to which G4-stabilization leads to low V_0_ values. From these results, TMPyP4 is clearly categorized as a G4-ligand (with V_0_ = 47.1, 32.0, 14.0 and 1.7 s^−1^ for 1, 5, 10 and 20 mol. equiv., respectively), more efficient than BRACO19 (with V_0_ between 67.6 and 49.9 s^−1^). Conversely, the tetra-anionic porphyrins help hybridization (with V_0_ up 104.0 s^−1^), thus demonstrating that the charge of the porphyrins matters: while cationic porphyrins stabilize G4 particularly at elevated concentrations (with V_0_ = 1.7 and 40.1 s^−1^ at 20 mol. equiv. of TMPyP4 and TEGPy, respectively), the negatively charged porphyrins improve the kinetics of the reaction at low (with V_0_ = 104.0 and 81.9 s^−1^ at 1 mol. equiv. of TPPS and TArPS, respectively) and elevated concentration, albeit less effectively (with V_0_ = 63.7 and 66.3 s^−1^ at 20 mol. equiv. of TPPS and TArPS, respectively). The other candidates moderately accelerate the hybridization (with V_0_ between 65.0-81.0, 60.5-87.1, 53.9-78.0, 56.9-70.7 and 65.7-73.6 s^−1^ for TEGP, Terpy, 1,5-BisNPO, 2,7-BisNPN and guaPhpC, respectively); PhpC was found quite active over the whole concentration range (with V_0_ between 72.7-94.5 s^−1^).

### CD titrations, UV-Vis spectroscopy, PAGE and FRET-melting investigations

We therefore decided to further investigate the properties of a panel of selected compounds, *i.e.*, TMPyP4 and PhenDC (likely G4-stabilizers) and TPPS, 1,5-BisNPO, 2,7-BisNPN and PhpC (likely G4-destabilizers), *via* a series of *in vitro* assays previously conducted to characterize possible G4-unwinding agents, *i.e.*, CD and PAGE. CD titrations were undertaken using the human telomeric G4-forming sequence (hTelo) and increasing amounts of candidates (1 to 10 mol. equiv.). Importantly, CD titrations were systematically paralleled with UV-Vis measurements, to investigate the spectroscopic behavior of both the small molecule and its complex with hTelo in solution. As seen in Figures 4A-C,G and S9-17, we first confirmed the previous observations according to which TMPyP4 triggers a strong decrease (68.7%, at 10 mol. equiv.) of the CD signal of the G4 (collected at its maximum, 293 nm). However, the UV-Vis contribution of TMPyP4 alone (Figure 4B, blue dotted line, and Figures 4C,G) where the G4 absorbs light (collected at its maximum, 257 nm) is important and dose-dependent, which also leads to an increase in the UV-Vis contribution of the TMPyP4/hTelo complex (from 3.5 to 36.5% variation, Figure 4B, blue line, and Figures 4C,G), implying a possible induced CD (iCD) contribution to the CD signatures of the TMPyP4/hTelo complex. PhenDC3 does not disrupt the G4 structure (2.5% variation) while its UV-Vis signatures are comparable to that of TMPyP4 (from-1.7 to 22.1% variation), implying again a possible iCD contribution. The UV-Vis contribution of both TPPS and PhpC, alone or in complex with the G4, are comparatively low (−4.1 to 13.0% for TPPS/hTelo, −3.4 to 7.7% for PhpC/hTelo) while they trigger significant CD decrease (down to −20.5 and −17.5%, respectively). The two azacyclophanes are found to trigger both CD (27.3 and 52.1% for 1,5-BisNPO and 2,7-BisNPN, respectively) and UV-Vis decreases (−26.7 and −20.5%, respectively), with a minimal UV-Vis contribution alone in solution. Collectively, these results highlight first and foremost that great caution must be exercised when relying only on CD titrations to study DNA/small molecule interactions, due to possible iCD contributions and other possibilities (*e.g*., aggregation, *vide infra*) that cannot be easily unraveled. Further investigations are thus required to better decipher the G4-interacting properties of these candidates.

**Figure 4.**
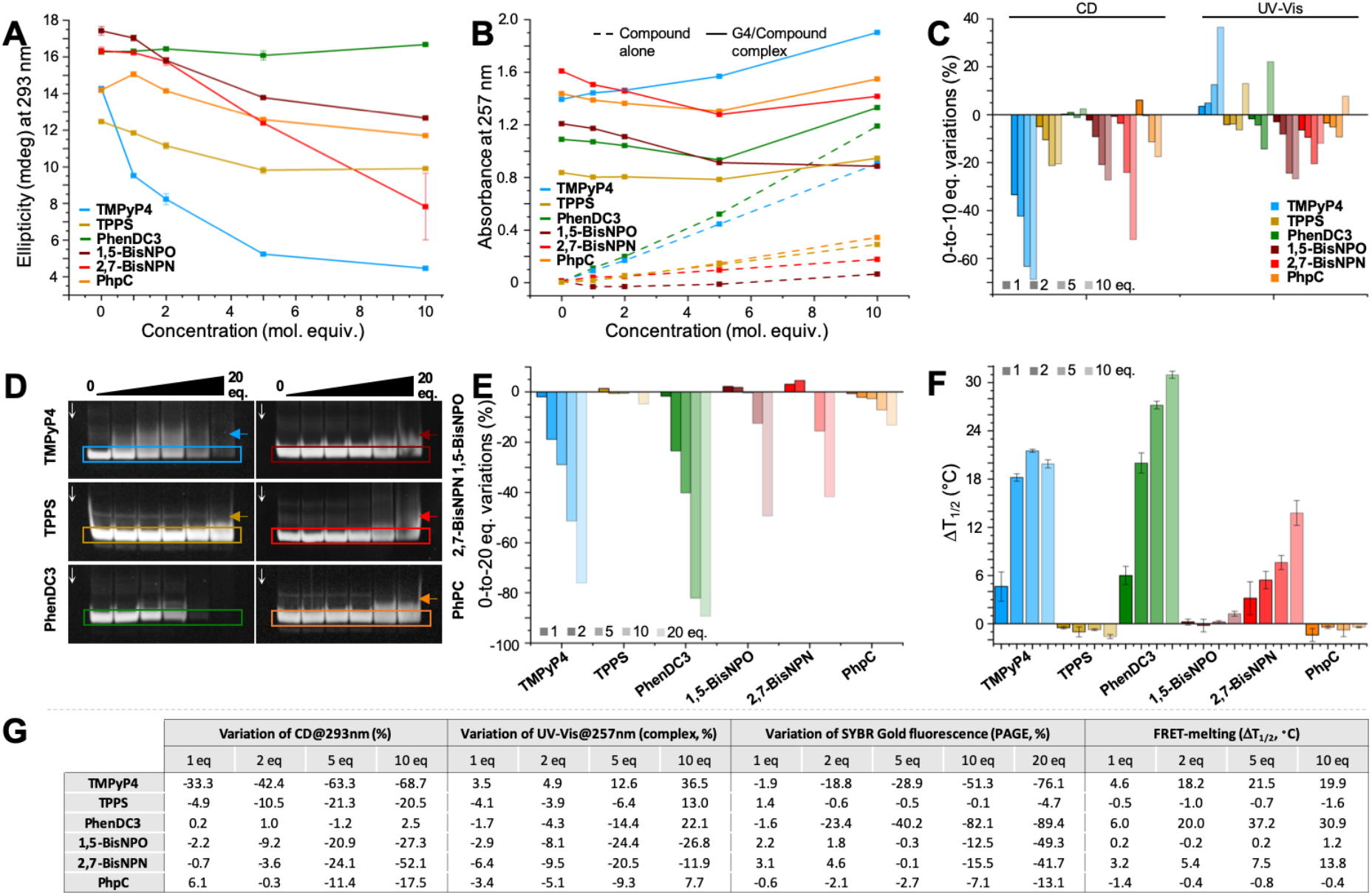
CD (**A**) and UV-Vis (**B**) titrations of hTelo G4 (3 μM) with increasing amounts (0 – 10 mol. equiv.) of 6 compounds (TMPyP4, TPPS, PhenDC3, 1,5-BisNPO, 2,7-BisNPN and PhpC) and summary (**C**) of the modifications of CD and UV-Vis signals (at 293 and 257 nm, respectively). PAGE results (**D**, n =2) and quantification (**E**) of experiments performed with hTelo and increasing amounts (0 – 20 mol. equiv.) of the 6 compounds. FRET-melting results (**F**, n = 3) of experiments performed with F21T and increasing amounts (0 – 10 mol. equiv.) of the 6 compounds. **G**. Summary of the variations observed in the CD and UV-Vis titrations, PAGE and FRET-melting assays performed with increasing amounts of the 6 compounds.

To do so, PAGE investigations were performed with this series of 6 compounds. In these conditions, the partial unfolding of hTelo G4 is expected to result in smeared PAGE bands (originating in an unstructured shape, a bigger molecular volume and a modified charge) rather than in loss of the signal. As above, TMPyP4 triggers a strong decrease of the band corresponding to hTelo (−76.1%, at 20 mol. equiv., Figures 4D,G), which is not in line with the UV-Vis titration (36.5% increase at 10 mol. equiv., Figures 4B,G) and might originate in possible aggregation/precipitation events. PhenDC3 leads to band disappearance to an even greater extent (−89.4%, at 20 mol. equiv.), again suggestive of possible aggregation/precipitation of the ligand/hTelo complex. Indeed, a ligand-mediated formation of multimeric G4s, or multimerization,^[52]^ is possible as it has been demonstrated for some G4-ligands (*e.g.*, *N*-methyl-indoloquinolinium^[53]^ and porphyrin)^[54]^ and characterized both experimentally^[55]^ and theoretically,^[56]^ which can lead to supramolecular assemblies too large to migrate within the gel lattice. In these conditions, TPPS is found rather inactive (from 1.4 to −4.7% variation) while the two azacyclophanes and PhpC provide dose-dependent responses (2.2 to −49.3% for 1,5-BisNPO, 3.1 to −41.7% for 2,7-BisNPN, −0.6% to - 13.1% for PhpC), in line with the CD/UV-Vis results. Therefore, and again, PAGE provides interesting insights into the G4-interacting properties of these candidates but cannot be used as a standalone technique since it is not devoid of experimental pitfalls.

Finally, we decided to evaluate the apparent affinity of these candidates for hTelo using the classical FRET-melting assay (with the doubly labelled hTelo, F21T). As seen in Figures 4F,G and S18-20, this stabilization is quite high and dose-dependent for PhenDC3 and 2,7-BisNPN (ΔT_1/2_ up to 30.9 and 13.8 °C, respectively, Figure 4G) while the saturation is obtained for 5 mol. equiv. of TMPyP4 (ΔT_1/2_ = 19.9 °C). Conversely, TPPS, 1,5-BisNPO and PhpC do not display any affinity for F21T and are even able to lower its melting temperature by 1.6, 0.2 and 1.4 °C, respectively. These results thus show that 3 candidates display high affinity for folded G4s (TMPyP4, PhenDC3 and 2,7-BisNPN) while TPPS, 1,5-BisNPO and PhpC do not interact with folded G4s. Collectively, the data gathered through this *in vitro* workflow indicate that only PhpC responded positively, TPPS failing at the PAGE step, 1,5-BisNPO at the CD/UV-Vis step and 2,7-BisNPN at the FRET-melting step. We thus decided to further investigate the G4-disrupting properties of PhpC through additional experiments.

### PhpC favors helicase processivity presumably *via* G4 disruption

First, we tried to gain direct insights into the way PhpC interacts with G4s but NMR investigations were poorly conclusive (Figure S21) owing to the overall decrease of the NMR signals of hTelo rather than a clear NMR signal redistribution. We thus exploited the fluorescence properties of the PhpC analogues, which are sensitive to the proximity of nucleobases. Initially embedded in a PNA strand, PhpC allowed for monitoring its association with the targeted DNA strand through fluorescence quenching.^[51]^ When titrated against increasing concentrations of guanosine monophosphate (GMP, 1 to 5 mol. equiv., to mimic one flipping G per G4), the PhpC fluorescence is marginally affected (−7.0% at best), indicating that the formation of the PhpC:GMP base pair does not influence the spectroscopic properties of the cytosine derivative, whatever the ionic content of the buffer (from 1 to 100 mM K^+^, Figure 5A). When titrated against hTelo, the K^+^-content of the buffer matters: decreasing the G4 stability by decreasing the K^+^ concentration of the buffer (from 100 to 1 mM K^+^) triggers a notable decrease of the PhpC fluorescence (−25.1, −30.5 and −36.4% for 100, 10 and 1 mM K^+^, respectively). The relationship between G4 stability (quantified as T_1/2_ values determined by FRET-melting assay) and fluorescence quenching as a function of the K^+^-content is almost linear (R^2^ = 0.96, see inset in Figure 5A). The decrease of the PhpC fluorescence might be attributed to the transient opening of the external G-quartet (the external G-quartet breathes more easily in a less stable G4), enabling PhpC to trap a flipping G (schematically represented in Figure 5A, left), thus laying in close proximity of the remaining G-triad that can affect its fluorescence by contact quenching.

**Figure 5.**
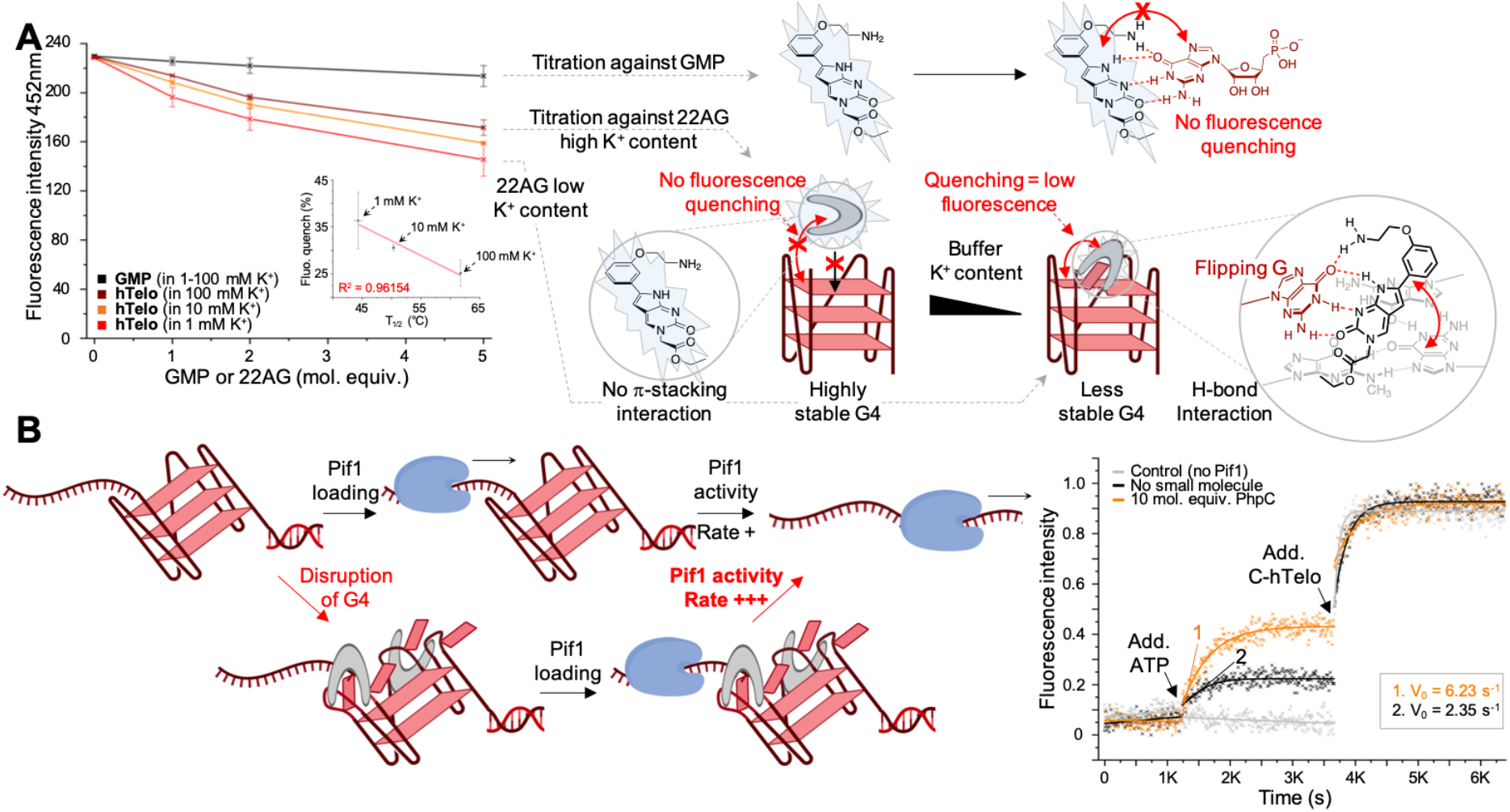
**A**. Fluorescence titration of PhpC (2 μM) upon increasing concentrations (1, 2 and 5 mol. equiv.) of GMP or 22AG, in buffer containing 1, 10 or 100 mM K^+^, as indicated (right panel; of note, the results collected with GMP were merged in a single curve); schematic representation of the hypotheses to explain observed fluorescence modifications (left panel). **B**. Schematic representation of the possible impact of the presence of a G4-disrupting molecule on the Pif1 processivity (right panel); results of the Pif1 helicase assay performed without Pif1 (control, grey), and with Pif1 in absence (black) or in presence of PhpC (10 mol. equiv.) (left panel).

We thus reasoned that this interaction might favor the G4 unfolding by Pif1, as a result of both an increase of the G4 instability by the transient stabilization of a partially open G4 (schematically represented in Figure 5B, right) and a weak and reversible interaction between PhpC and a wobbling G, which might be easily disrupted during Pif1 translocation. To investigate this, the complete Pif1 helicase assay (described in Figure 1A) was implemented with 2 enzyme concentrations (160 and 170 nM) in absence or presence of 10 mol. equiv. PhpC (Figures 5B, right, and S22). Quite satisfyingly, the presence of PhpC enhances the Pif1-mediated G4 unfolding (between 2.4- and 4.0-fold, Figure 5B, left, and S22), while all other small molecules evaluated so far were reported to impede it.^[37]^ These results thus open new horizons for chemical biology, as they show that small molecules can facilitate G4 unwinding by G4-helicases in a nature-inspired manner, given that only proteins such as RPA (replication protein A)^[57]^ have been reported to date to stimulate Pif1 activity. They also provide the first example of a small molecule able to do so, thus offering new strategic opportunities for compensating and/or rescuing the G4-helicase deficiencies underlying severe genetic dysfunctions and diseases.^[7]^

## Conclusion

The wealth of data collected here highlight the issues faced when exploring the ability of small molecules to disrupt G4s, as their behavior is found to be strongly dependent on the technique and the concentration used, as previously evoked.^[35]^ This originates in the fact that small molecules can interact with G4s in many different ways, as confirmed here with TMPyP4, certainly the most representative example of compound whose G4-stabilization/disruption properties are complicated to unravel. These results keep on demonstrating also the versatility of the porphyrins as DNA-interacting scaffolds as modification of their chemical core (here, their charges and side-arms; previously their side-arms^[35]^ and the presence of a metal in their central cavity)^[34]^ can reverse their binding properties. They also cast a bright light on the promising G-clamp analog scaffold PhpC to efficiently disrupt G4 structures and facilitate G4-helicase activity *in vitro*.

Beyond this, they lend credence to the reliability of the G4-unfold assay described here to detect putative G4 unwinders and, above all, to the step-by-step methodology relying on a combination of techniques (G4-unfold, CD, PAGE, FRET-melting, Pif1 helicase assay) to assess the actual efficiency of putative G4-unwinding candidates in the most reliable possible way. Applying this workflow to wider chemical libraries will undoubtedly lead to the identification of ever more efficient G4-unwinders, which will find soon applications as promising chemical biology tools in the field of genetic diseases.

## Materials and Methods

### Oligonucleotides

All oligonucleotides used here were purchased from Eurogentec (Seraing, Belgium) and stored at −20°C as 500 μM stock solutions in deionized water (18.2 MΩ.cm resistivity). The actual concentration of these stock solutions was determined through a dilution to 5 μM theoretical concentration *via* a UV spectral analysis at 260 nm after 5 min at 90 °C. Quadruplex structures of hTelo (for CD and PAGE investigations) were prepared by mixing 10 μL of the constitutive strand (500 μM) with 10 μL of a lithium cacodylate buffer solution (100 mM, pH 7.2), plus 10 μL of a KCl/LiCl solution (100 mM/900 mM) and 70 μL of Milli-Q water. Quadruplex structures of F21T (for FRET-melting) were prepared by mixing 5 μL of the constitutive strand (500 μM) with 10 μL of a lithium cacodylate buffer solution (100 mM, pH 7.2), plus 10 μL of a KCl/LiCl solution (100 mM/900 mM) and 75 μL of Milli-Q water. The final concentrations were theoretically 25 μM (F21T) and 50 μM (hTelo). The DNA system required for the G4-unfold and the Pif1 helicase assay was annealed at 1 μM concentration by mixing 1 μM Dabcyl-labeled oligonucleotide and 0.85 μM FAM-labeled oligonucleotide in 20 mM Tris-HCl buffer (pH 7.2, 5 mM MgCl_2_, 1 mM KCl and 99 mM NaCl). All higher-order structures were folded *via* a short heating step (90 °C, 5 min) followed by gradual cooling steps (80, 60, 50, 40 and 30 °C (10 min for 80°C step and 1 h/step for others), 25 °C (2 h)) prior to be stored at 4 °C overnight before use and/or stored at −20 °C.

### The G4-unfold assay

Every reactions were carried out in duplicate in 96-well plates (Greiner Bio-one; 96-well, black, flat bottom) at 25 °C and fluorescence monitored in a microplate reader (BMG Labtech ClarioStar). To a 10-μL solution of 40 nM S-hTelo in 20 mM Tris-HCl buffer pH 7.2 (5 mM MgCl_2_, 1 mM KCl and 99 mM NaCl for experiments without molecule (control) or 10 mM MgCl_2_, 1 mM KCl and 99 mM NaCl for experiments performed in the presence of molecules) were added 40 μL of candidates in Tris-HCl buffer (0 + 40 μL for the control; 1 + 39 μL for 1 mol. equiv.; 5 + 35 μL for 5 mol. equiv.; 10 + 30 μL for 10 mol. equiv.; 20 + 20 μL for 20 mol. equiv.) and the fluorescence emission was recorded every 10 s (λ_ex_ = 492 nm; λ_em_ = 516 nm) till the emission signal was stable (15 min). Next, 2.2 μL (2 μM, 5 mol. equiv.) of C-htelo was added to every well, the plates were stirred for 10 s, and emission was monitored every 10 s during 25 min. Final data were analyzed with Excel (Microsoft Corp.) and OriginPro®9.1 (OriginLab Corp.).

### CD and UV-Vis titrations

CD and UV-Vis spectra were recorded on a JASCO J-815 spectropolarimeter in a 10 mm path-length quartz semi-micro cuvette (Starna). CD spectra were recorded over a range of 210-350nm (bandwidth = 1 nm, 1 nm data pitch, 1 s response, scan speed = 200 nm.min^−1^, averaged over 4 scans, zeroed at 350 nm). Samples were prepared in 100 μL (final volume) comprising 6 μL hTelo (3 μM final concentration) in 10 mM lithium cacodylate buffer (pH 7.2), 10 mM KCl and 90 mM LiCl without and with increasing amounts of candidates (3, 6, 15 and 30 μM in H_2_O or DMSO; 3 μL molecule (100 μM) for 1 mol. equiv. (twice) then 11 μL molecule (100 μM) for 5 mol. equiv. and 1.7 μL molecule (1 mM) for 10 mol. equiv.). Final data were analyzed with Excel (Microsoft Corp.) and OriginPro®9.1 (OriginLab Corp.).

### PAGE analyses

Nondenaturing polyacrylamide gel electrophoresis was carried out with 20% polyacrylamide-bisacrylamide gel. Samples were prepared in 15 μL (final volume) comprising 1.5 μL DNA (5 μM final concentration) and increasing amounts of candidates (5, 10, 25, 50 and 100 μM in H_2_O or DMSO; 0 μL + 11 μL of 10mM lithium cacodylate buffer (pH 7.2), 10 mM KCl and 90 mM LiCl for the control; 0.75 μL molecule (100 μM) + 10.25 μL buffer for 1 mol. equiv.; 1.5 μL molecule (100 μM) + 9.5 μL buffer for 2 mol. equiv.; 3.75 μL molecule (100 μM) + 7.25 μL buffer for 5 mol. equiv.; 0.75 μL molecule (1 mM) + 10.25 μL buffer for 10 mol. equiv.; 1.5 μL molecule (1 mM) + 9.5 μL buffer for 20 mol. equiv.) and 2.5 μL 6X DNA loading buffer (Thermo Fisher Scientific). 10 μL of this mixture were thus loaded on the gel. The electrophoretic migration was performed in 1×TBE (tris-borate EDTA buffer), pH 8.3, for 15 min at 7 W then 35 min at 12 W, at 4°C. After the migration, gels were analyzed after a post-staining step (SYBR®Gold solution, 1:133 (10 μL in 75 mL of 1×TBE), 15 min, 25°C under gentle agitation) with a UVP MultiDoc-It® imaging system (λ_ex_ = 254 nm).

### FRET-melting assay

Experiments were performed in a 96-well format using a Mx3005P qPCR machine (Agilent) equipped with FAM filters (λ_ex_ = 492 nm; λ_em_ = 516 nm) in 100 μL (final volume) of 10 mM lithium cacodylate buffer (pH 7.2) plus 10 mM KCl/90 mM LiCl with 0.2 μM of F21T and 0.2, 0.4, 1.0 or 2.0 μM of both candidates. After a first equilibration step (25 °C, 30 s), a stepwise increase of 1 °C every 30 s for 65 cycles to reach 90 °C was performed, and measurements were made after each cycle. Final data were analyzed with Excel (Microsoft Corp.) and OriginPro®9.1 (OriginLab Corp.). The emission of FAM was normalized (0 to 1), and T_1/2_ was defined as the temperature for which the normalized emission is 0.5; ΔT_1/2_ values, calculated as follows: ΔT_1/2_ = [T_1/2_(F21T+ligand)-T_1/2_(F21T)], are means of 3 experiments.

### Pif1 helicase assay

According to O. Mendoza *et al*.^[37]^ Helicase reactions were carried out in duplicate in 96-well plates (Greiner Bio-one; 96-well, black, flat bottom) at 25 °C and fluorescence monitored in a microplate reader (BMG Labtech ClarioStar). Every well contained a 50-μL solution of 40 nM S-htelo in 20 mM Tris-HCl buffer (5 mM MgCl_2_, 1 mM KCl and 99 mM NaCl for experiments without molecule (control) or 10 mM MgCl_2_, 1 mM KCl and 99 mM NaCl for experiments performed in the presence of molecules), 160 or 170 nM of Pif1 enzyme and 200 nM of Trap oligonucleotide (unlabeled, complementary to the FAM-labelled strand). Next, 1 μL of PhpC (20 μM, 400 nM, 10 mol. equiv.) or Tris-HCl buffer (control) and the fluorescence emission was recorded every 10 s (λ_ex_ = 492 nm; λ_em_ = 516 nm) till the emission signal was stable (15 min). Next, 5 μL of a 50 mM ATP solution was added to every well, the 96-well plate was stirred for 10 s and the fluorescence emission was recorded every 10 s till the emission signal was stable (20 min). Finally, 5 μL (2 μM, 10 mol. equiv.) of C-htelo was added to every well, the plates were stirred for 10 s, and emission was monitored every 10 s during 25 min. Final data were analyzed with Excel (Microsoft Corp.) and OriginPro®9.1 (OriginLab Corp.).

## Acknowledgments

The authors thank the Centre National de la Recherche Scientifique (CNRS), the Agence Nationale de la Recherche (ANR-17-CE17-0010-01), the Université de Bourgogne, Conseil Régional de Bourgogne and the European Union (PO FEDER-FSE Bourgogne 2014/2020 programs) and the INSERM Plan Cancer 2014-2019 (n° 19CP117-00) for financial support.

## Abbreviations

^§^ TCPP stand for: meso-tetra(4-carboxyphenyl)porphine, also known as 4,4’,4’’,4’’’-(porphine-5,10,15,20-tetrayl)tetrakis(benzoic acid) or meso-Tetraphenylporphine-4,4’,4’’,4’’’-tetracarboxylic acid (Spm for spermine);
TMPyP4 for: 5,10,15,20-tetrakis-[N-methyl-4-pyridyl]-21H,23H-porphyrin
TEGPY for: tetra-[(ethylene glycol)-pyridyl]-porphyrin, also known as 5,10,15,20-tetrakis-[N-(2-(2-(2-methoxy)-ethoxy)-ethoxy)-ethyl-4-pyridyl]-21H,23H-porphyrin;
TEGP for: tetra-[ethylene glycol]-porphyrin, also known as 5,10,15,20-tetrakis-[4-(2-(2-(2-methoxy)-ethoxy)-ethoxy)-ethyl]-21H,23H-porphyrin
TArPS for: 5,10,15,20-tetrakis[3-sulfonato-4-O-[2-[2-(2-methoxy)ethoxy]ethoxy]ethylphenyl]-21H,23H-porphyrin
TPPS for: 5,10,15,20-tetrakis[4-sulfonatophenyl]-21H,23H-porphyrin
PhpC for: 6-phenylpyrrolocytosine (gua for guanidine).

---

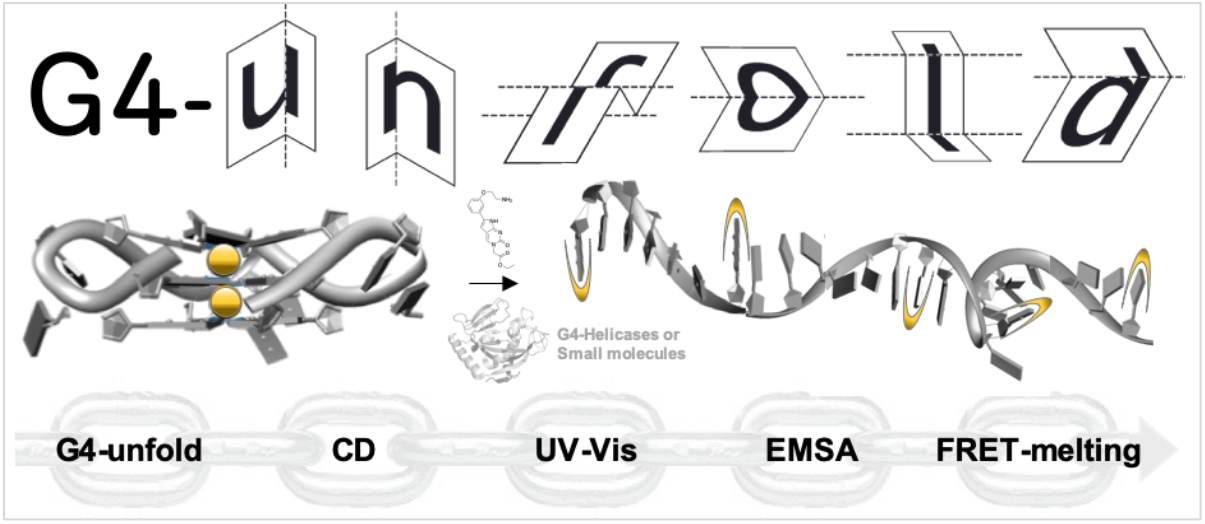
Graphical abstract.

