## Supplementary Information for "Disclosing the actual efficiency of G-quadruplex-DNA–disrupting small molecules"

### -- Supporting Information --

#### 1. G4-unfold assay

The detailed protocol is described in the main manuscript.

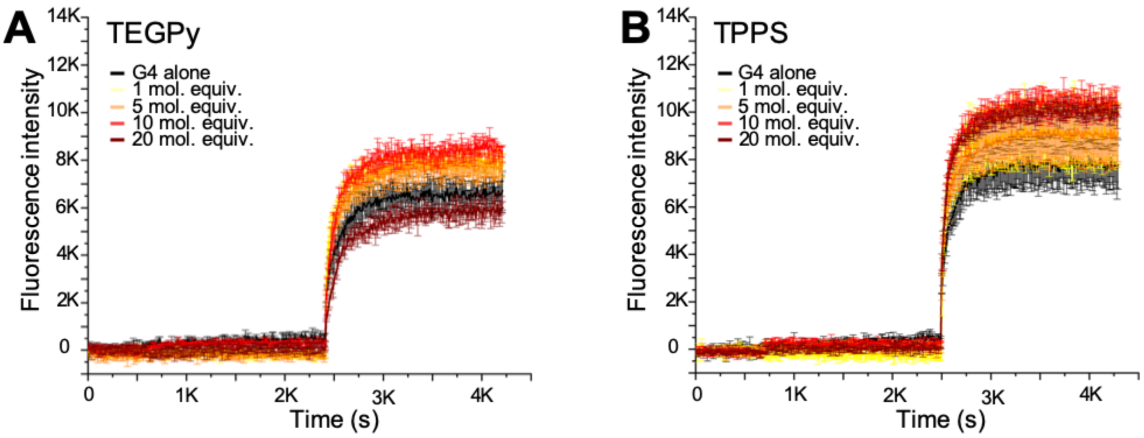


**Figure S1**. G4-unfold results collected with TEGPy and TPPS

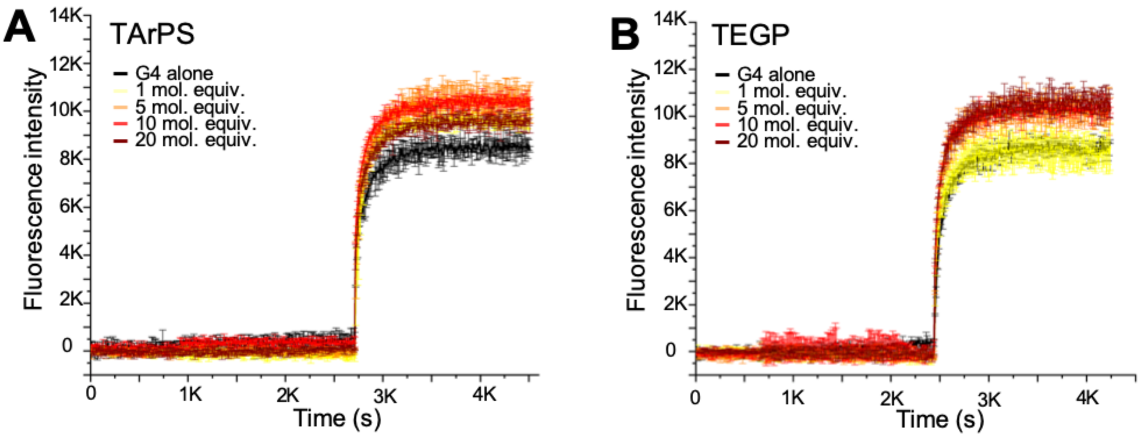


**Figure S2**. G4-unfold results collected with TArPS and TEGP

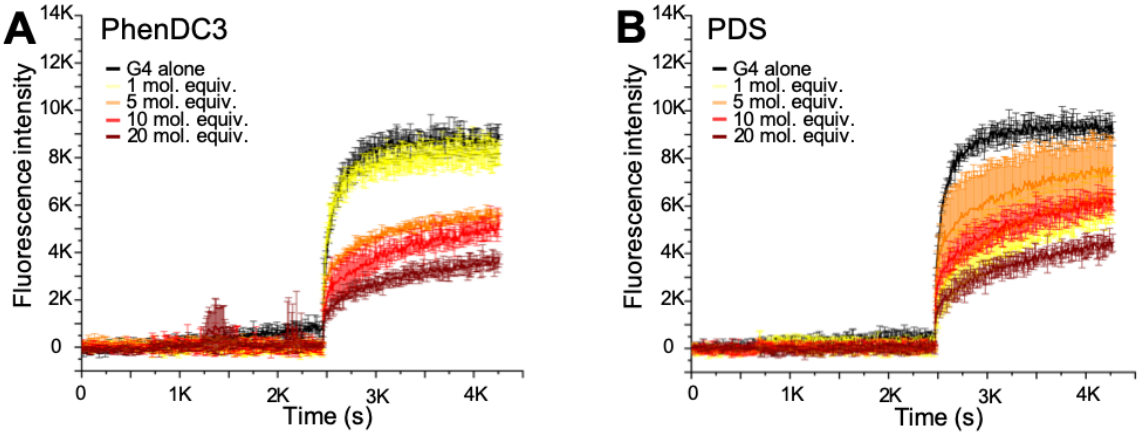


**Figure S3**. G4-unfold results collected with PhenDC3 and PDS

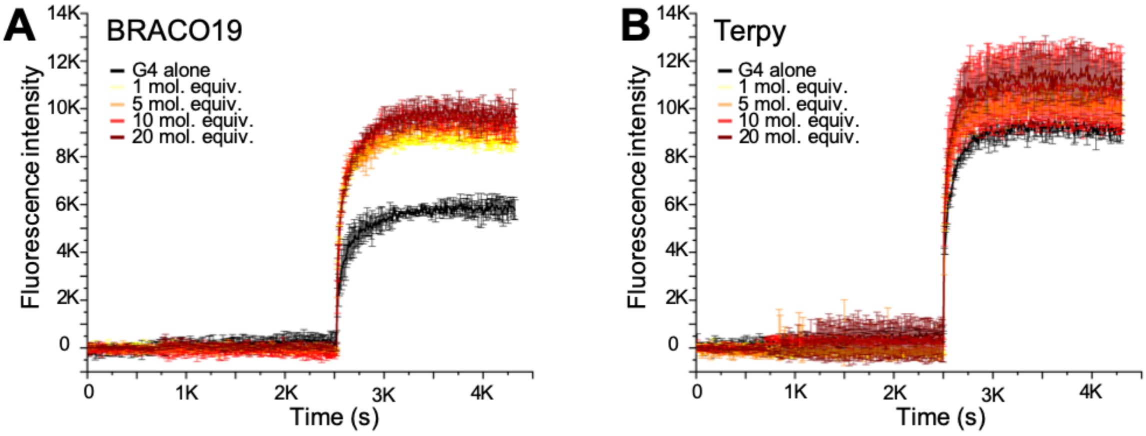


**Figure S4**. G4-unfold results collected with BRACO19 and Terpy

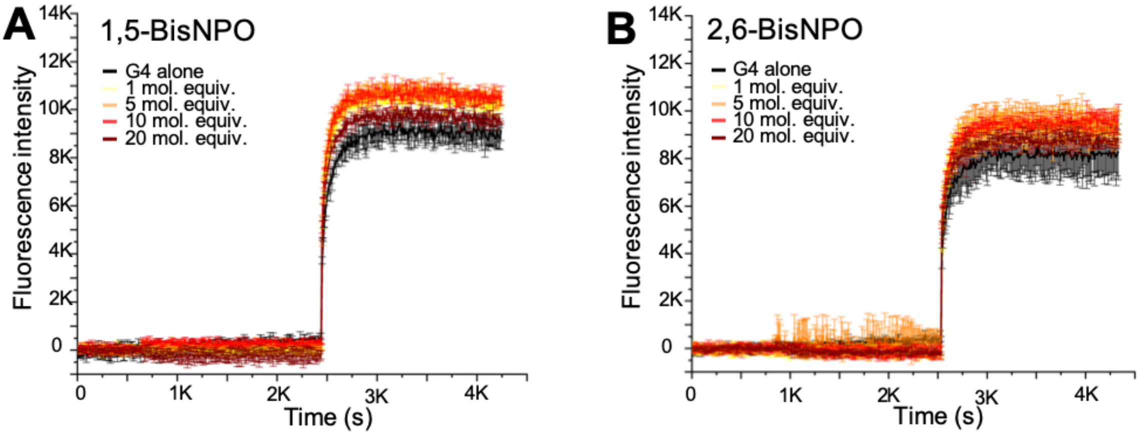


**Figure S5**. G4-unfold results collected with 1,5-BisNPO and 2,6-BisNPO

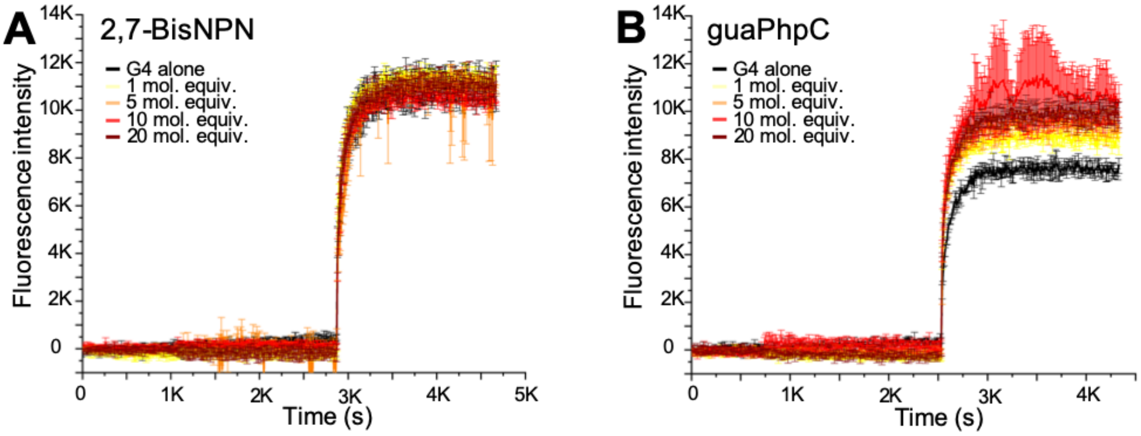


**Figure S6**. G4-unfold results collected with 2,7-BisNPN and guaPhpC

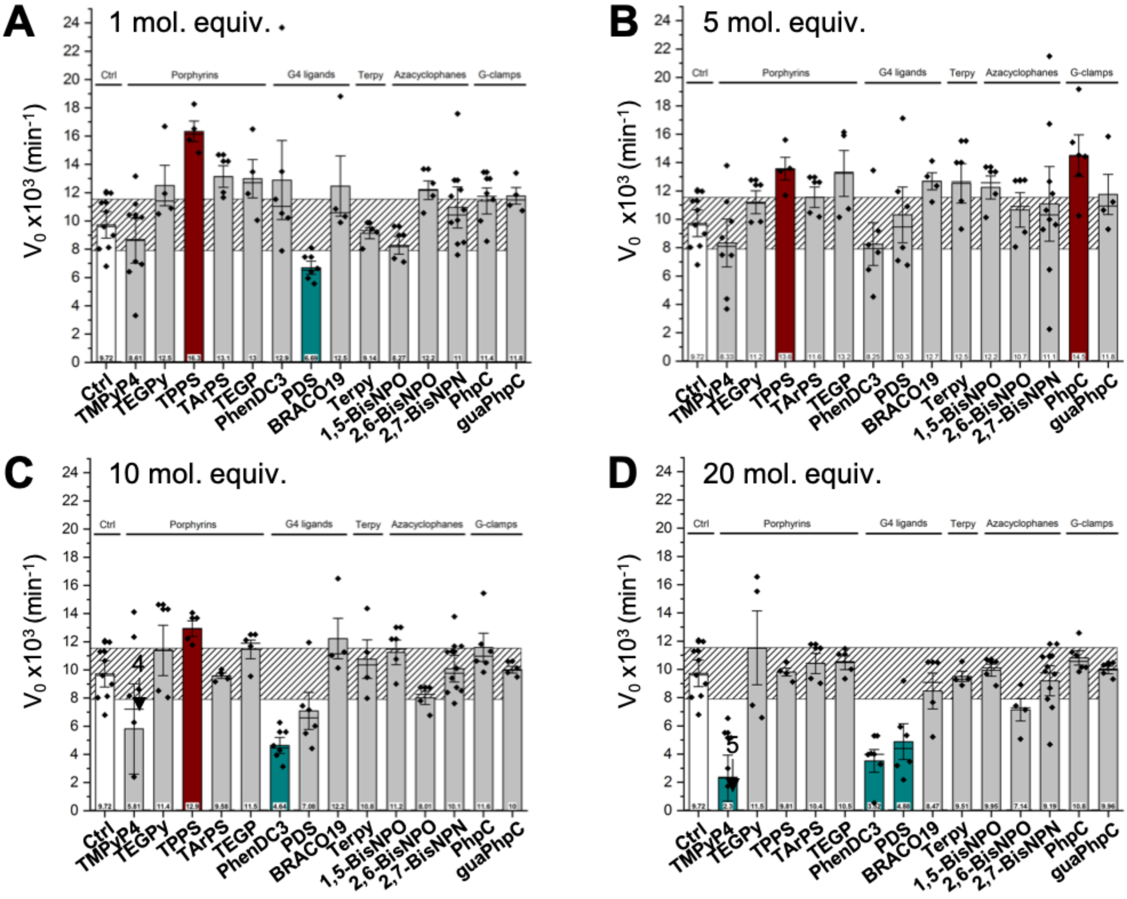


**Figure S7. A-D**. Normalized results ($V_0$, expressed in $s^{-1}$) of the G4-unfold assay performed with the FAM/dabcyl duplex construct (40 nM) and increasing amounts of 14 compounds (from 1 to 20 mol. equiv.); diamonds are the experimental data (n > 4); bars represent the averaged $V_0$; error bars represent standard deviation (S.D.); the diagonally hashed grey zone represent the exclusion zone (calculated as 2xS.D. of the control, Ctrl).

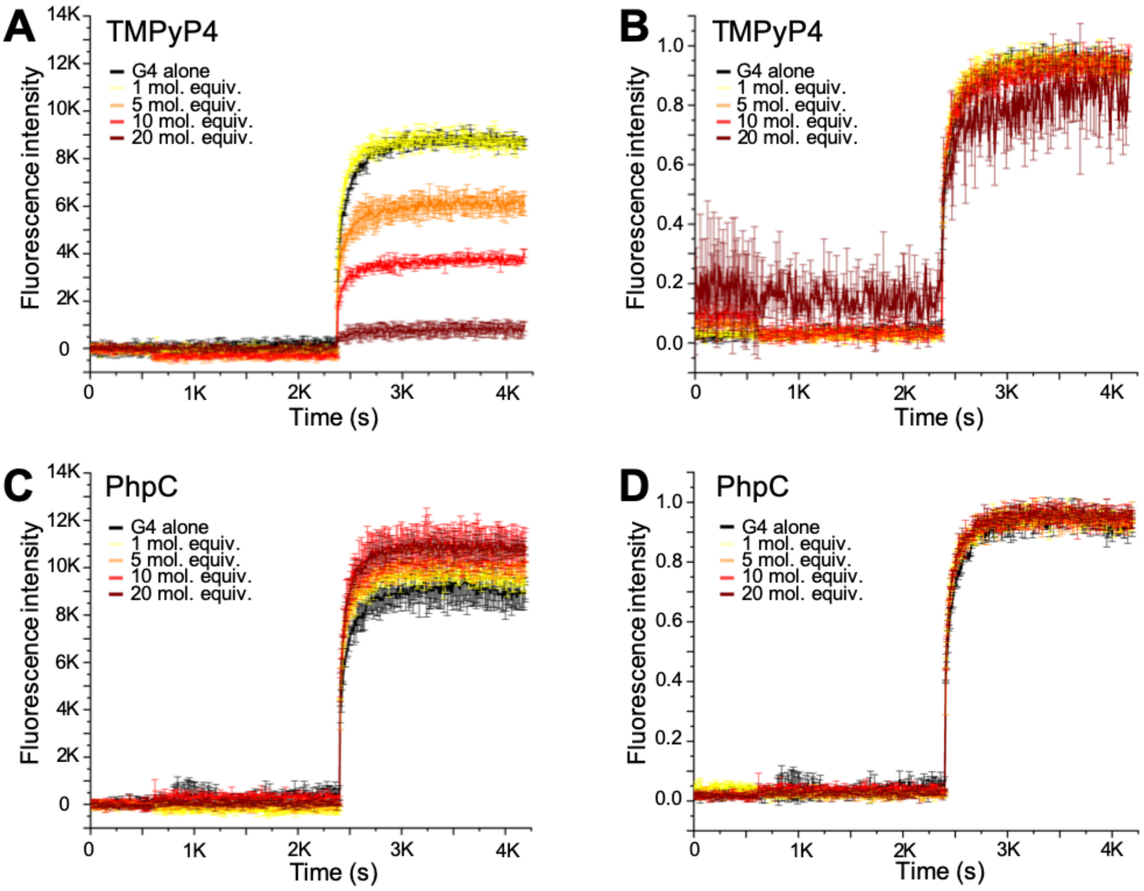


**Figure S8. A-D**. Comparison of the raw (A, C) and normalized curves (B, D) obtained for the G4-unfold assay performed with TMPyP4 (A, B) and PhpC (C, D).

### 2. CD titrations

The detailed protocol is described in the main manuscript.

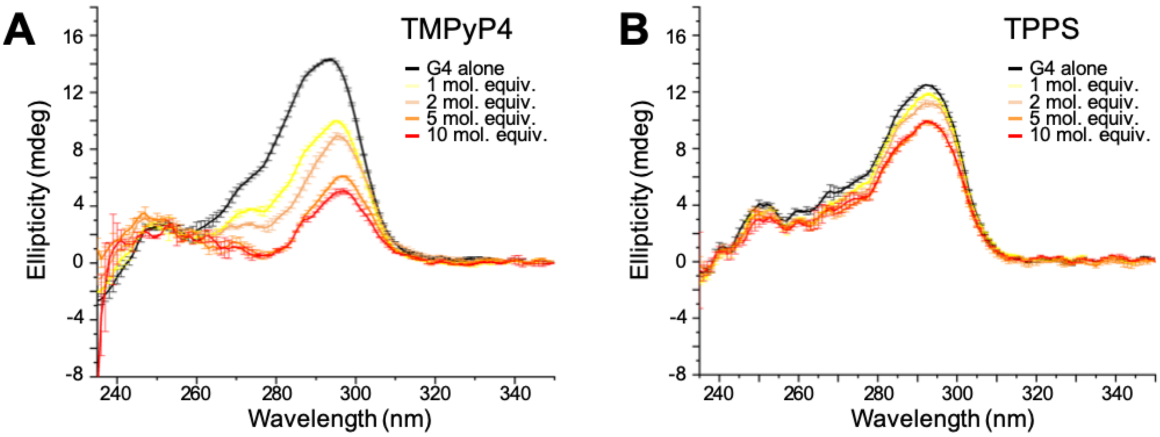


**Figure S9**. CD titrations of hTelo by increasing amounts of TMPyP4 and TPPS

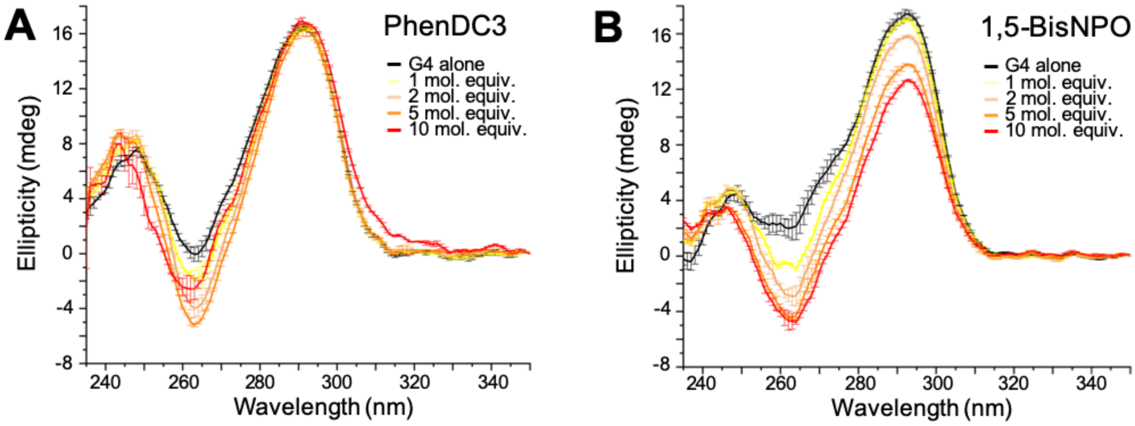


**Figure S10**. CD titrations of hTelo by increasing amounts of PhenDC3 and 1,5-BisNPO

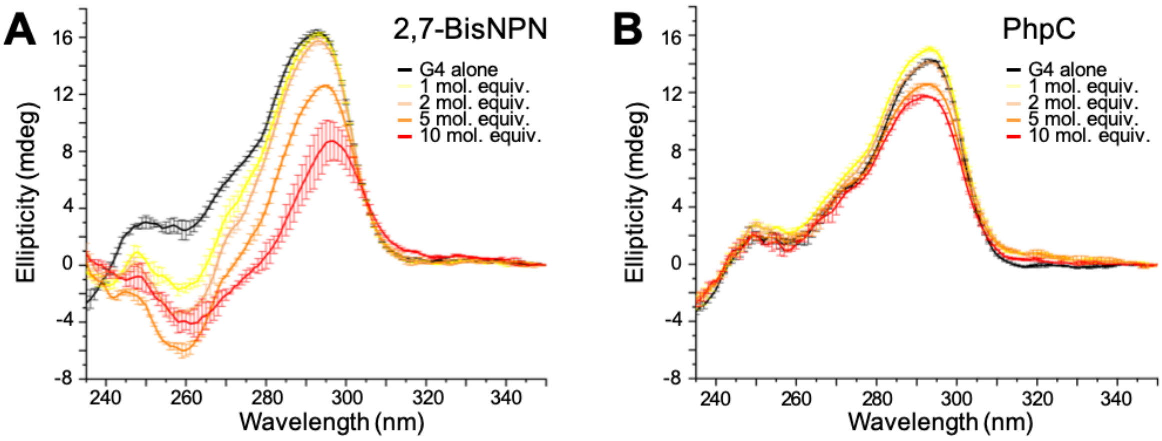


**Figure S11**. CD titrations of hTelo by increasing amounts of 2,7-BisNPN and PhpC

### 3. UV-Vis titrations

The detailed protocol is described in the main manuscript.

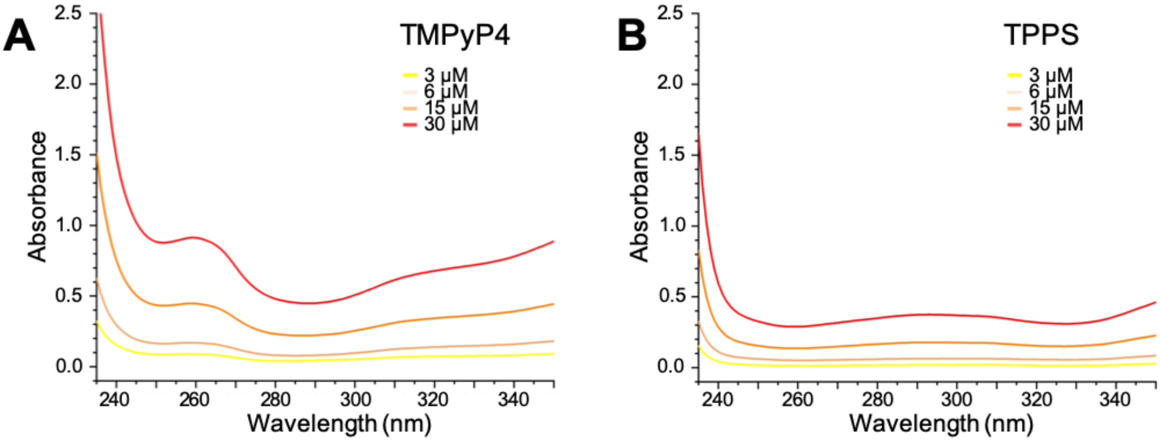


**Figure S12**. UV-Vis spectra of increasing amounts of TMPyP4 and TPPS

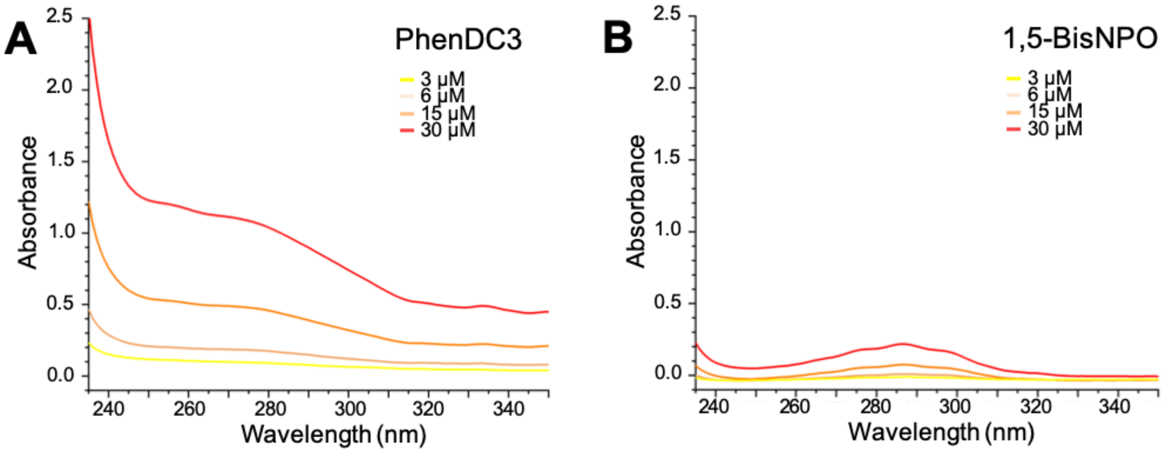


**Figure S13**. UV-Vis spectra of increasing amounts of PhenDC3 and 1,5-BisNPO

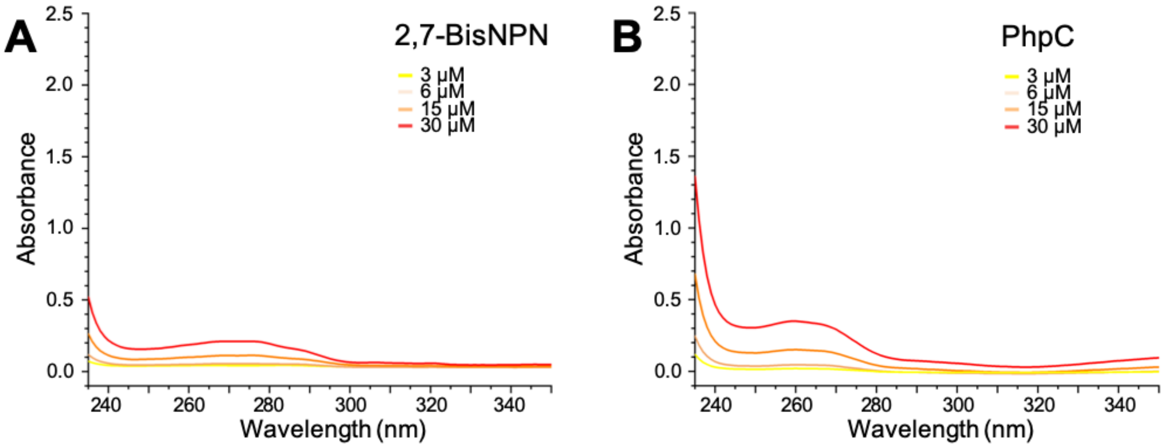


**Figure S14**. UV-Vis spectra of increasing amounts of 2,7-BisNPN and PhpC

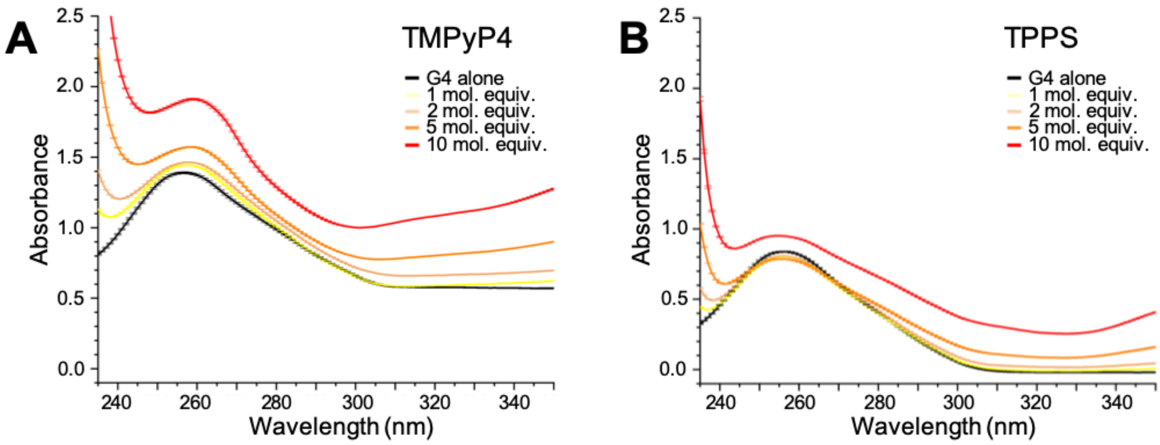


**Figure S15**. UV-Vis titrations of hTelo by increasing amounts of TMPyP4 and TPPS

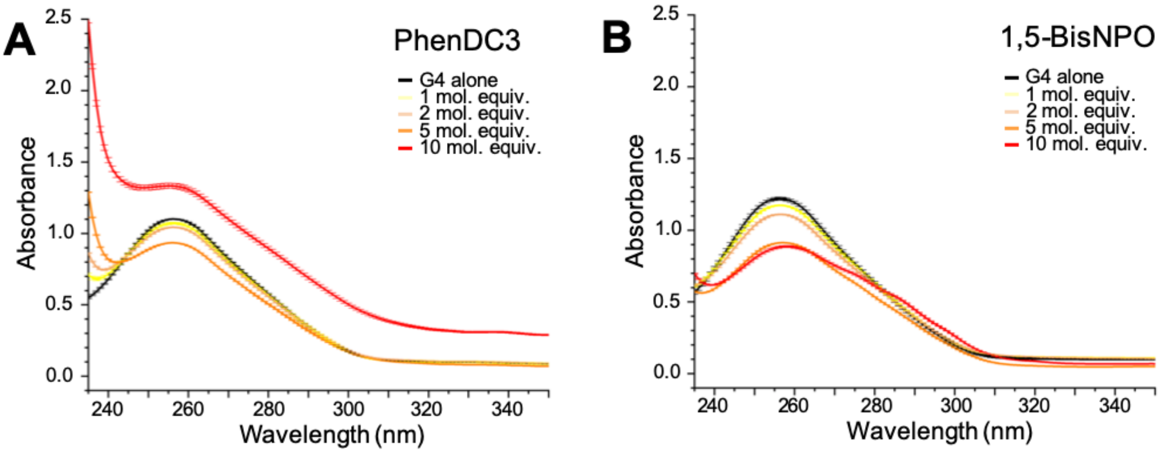


**Figure S16**. UV-Vis titrations of hTelo by increasing amounts of PhenDC3 and 1,5-BisNPO

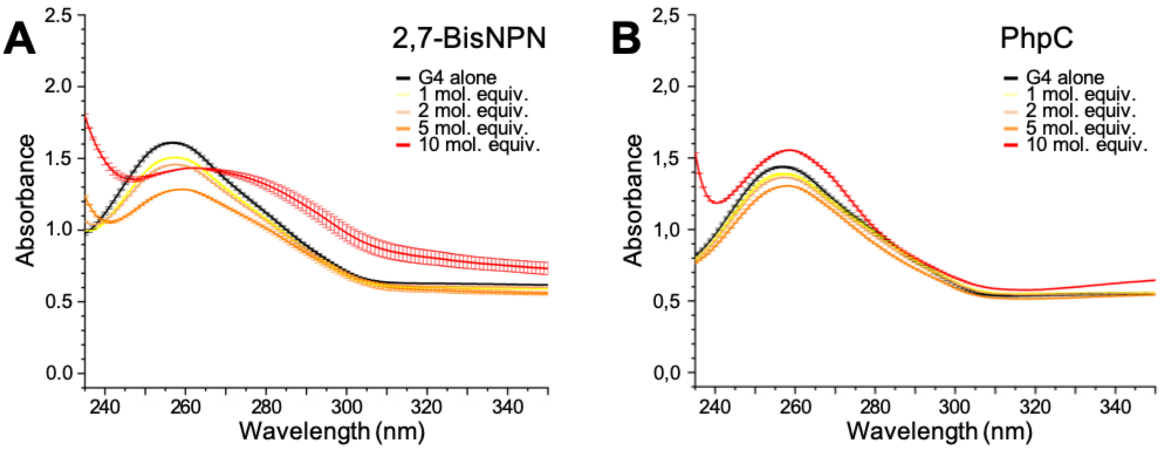


**Figure S17**. UV-Vis titrations of hTelo by increasing amounts of 2,7-BisNPN and PhpC

### 4. FRET-melting assay

The detailed protocol is described in the main manuscript.

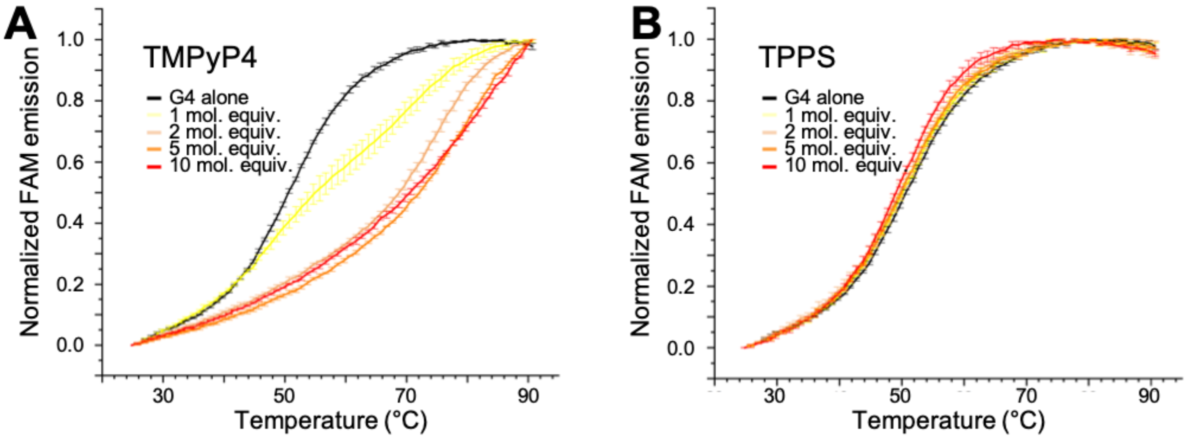


**Figure S18**. FRET-melting experiments performed with F21T by increasing amounts of TMPyP4 and TPPS

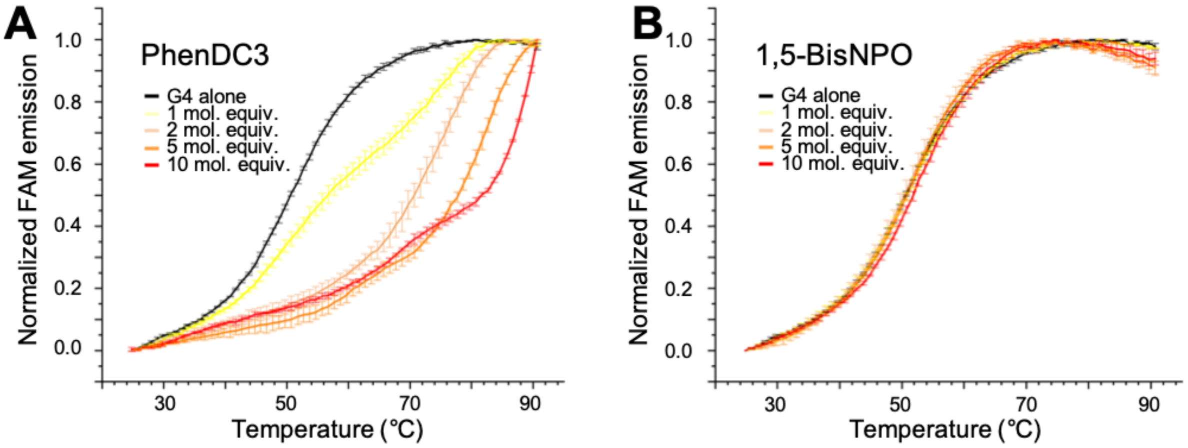


**Figure S19**. FRET-melting experiments performed with F21T by increasing amounts of PhenDC3 and 1,5-BisNPO

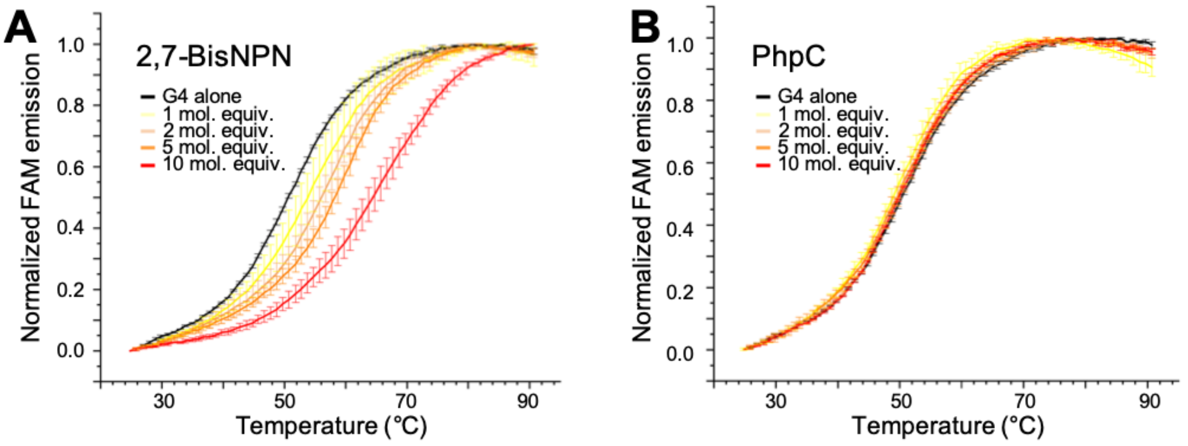


**Figure S20**. FRET-melting experiments performed with F21T by increasing amounts of 2,7-BisNPN and PhpC

### 5. NMR investigations

$^{1}H$ NMR spectra were acquired at 298K on a Bruker Avance II 600MHz spectrometer with Prodigy cryprobe 5mm BBOF using the “1D_exitation sculpting” solvent suppression sequence (zgesgp). $^{1}H$ NMR spectrum were obtained after 4248 scans in 2.5mm NMR tube. Samples were prepared in 250 µL comprising 200 µL hTelo in 10 mM lithium cacodylate buffer (pH 7.2), 10 mM KCl and 90 mM LiCl with 10% of $D_2O$ in presence of DSS (internal standard) without and with increasing amounts of PhpC in water (twice 25 µL of 10 mM solution). Final data were analyzed with Topspin 4.0.6.

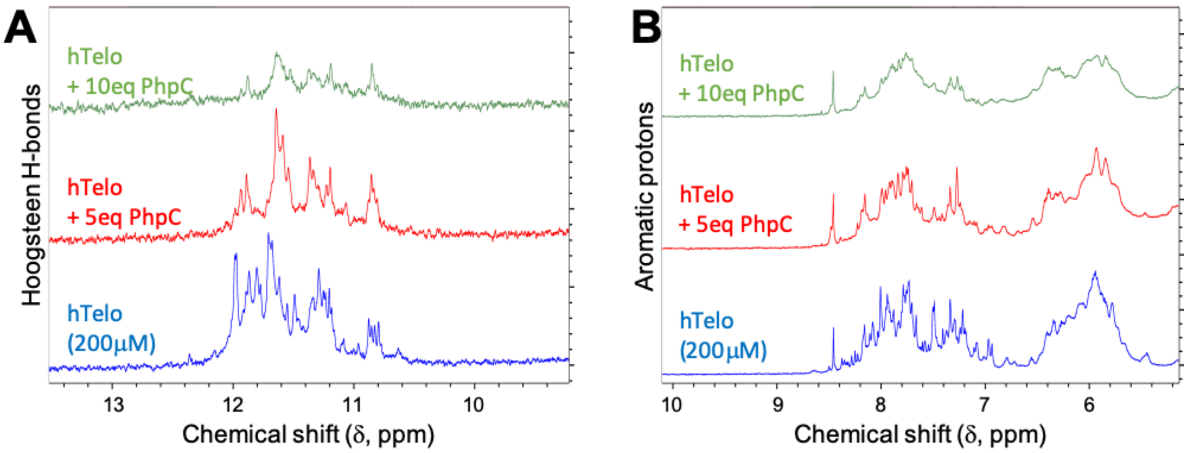


**Figure S21**. NMR titration (parts) of hTelo by increasing amounts of PhpC

### 6. Pif1 Helicase assay

The detailed protocol is described in the main manuscript.

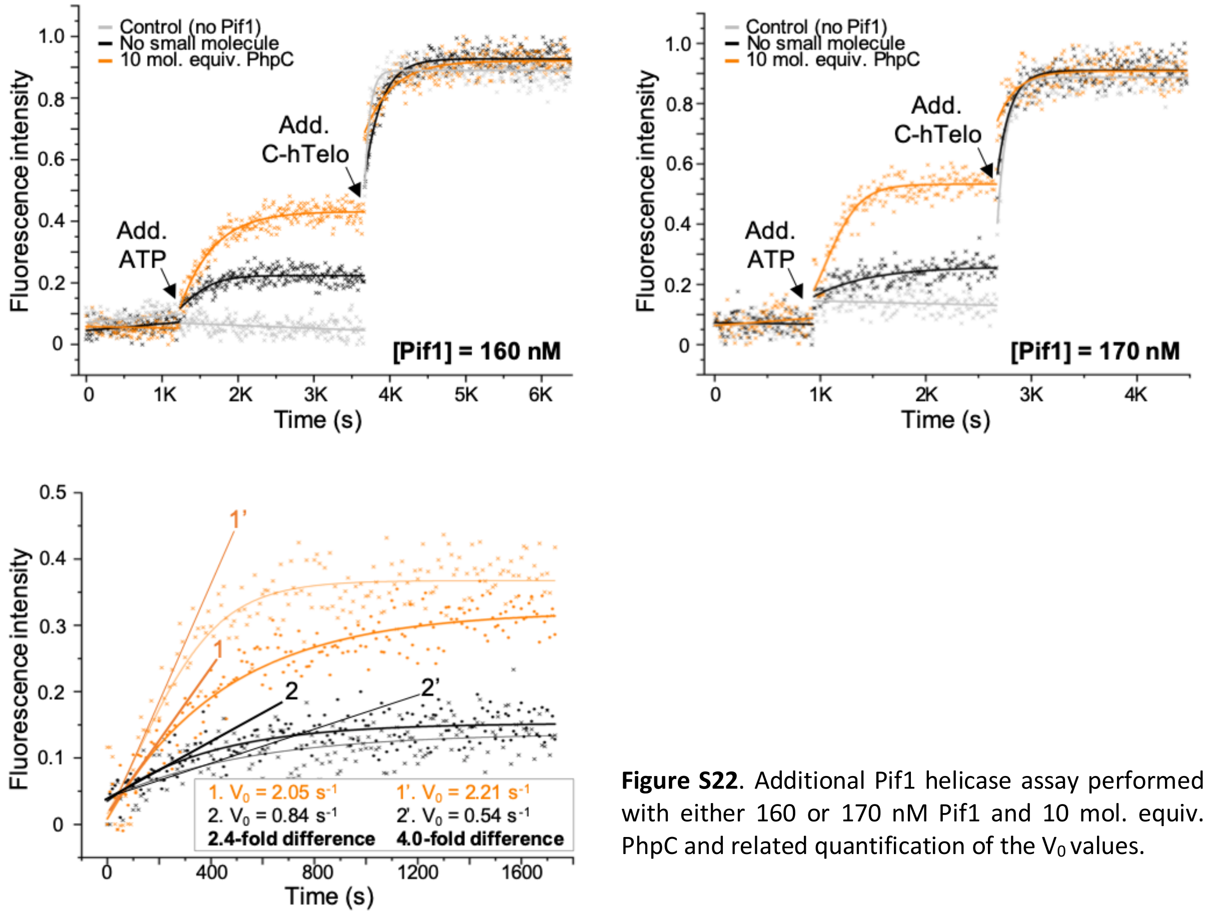


**Figure S22**. Additional Pif1 helicase assay performed with either 160 or 170 nM Pif1 and 10 mol. equiv. PhpC and related quantification of the $V_0$ values.
